# Topology of the U12-U6atac snRNA complex of the minor spliceosome and binding by NTC-related protein RBM22

**DOI:** 10.1101/825604

**Authors:** Joanna Ciavarella, William Perea, Nancy L. Greenbaum

## Abstract

Splicing of precursor messenger RNA is catalyzed by the spliceosome, a dynamic ribonucleoprotein assembly composed of five small nuclear (sn)RNAs and >100 proteins. RNA components catalyze the two transesterification reactions, but proteins perform critical roles in assembly and rearrangement. The catalytic core comprises a paired complex involving U2 and U6 snRNAs for the major form of the spliceosome and U12 and U6_atac_ snRNAs for the minor variant (~0.3% of all spliceosomes in higher eukaryotes); the latter performs identical chemistry, despite limited sequence conservation outside key catalytic elements, and lack of the multi-stem central junction found in the U2-U6 snRNA complex. Here we use solution NMR techniques to show that base pairing patterns of the U12-U6_atac_ snRNA complex of both human and *Arabidopsis* share key elements with the major spliceosome’s U2-U6 snRNA complex; probing of the single-stranded segment opposing termini of the snRNAs indicates elongation in this region in place of the stacked base pairs at the base of the U6 intramolecular stem loop in the U2-U6 snRNA complex. Binding affinity of RBM22, a protein implicated in remodeling human U2-U6 snRNA prior to catalysis, to U12-U6_atac_ was analyzed by electrophoretic mobility shift assays in which we monitored migration of both protein and RNA components in the same gel. Results indicate that RBM22 binds the U2-U6 and U12-U6_atac_ snRNA complexes specifically and with K_d_ = 3.5 µM and 8.2 µM, respectively. Similar affinity between RBM22 and each RNA complex suggests that the protein performs the same role in both spliceosomes.

## INTRODUCTION

The excision of noncoding intervening sequences (introns) and ligation of flanking coding regions (exons) from precursor messenger (pre-m)RNA is mediated by the spliceosome, a large and dynamic ribonucleoprotein complex found in the nuclei of eukaryotic cells. The vast majority of spliceosomes comprise the snRNAs U1, U2, U4, U5, and U6 in association with at least 150 proteins (actual number depends upon the organism and state of assembly) to form snRNPs (rev. by Will and Lührmann 2011;Papasaikas and Válcarcel 2016). Pre-mRNA splicing is a critical step in the maturation of pre-mRNA into translatable mRNA transcripts and involves two sequential transesterification reactions catalyzed by RNA components of the spliceosome; *i.e.* the spliceosome is a ribozyme (Fica et al. 2014).

During the first transesterification reaction, the 2’OH of a conserved adenosine residue in the intron, referred to as the branch site due to the branched lariat intermediate it forms, performs a nucleophilic attack at the 5’ splice site, forming a free 3’OH on the exon (rev. by Moore et al. 1993). In the second reaction, the free 3’OH undergoes an attack at the 3’ splice site, joining the two exons while releasing the lariat intron. These two transesterification reactions have been previously proposed to involve a two-metal ion center where one Mg^2+^ ion activates the nucleophilic 2’OH while the second Mg^2+^ ion stabilizes the oxyanion leaving group (Steitz and Steitz, 1993).

Throughout the splicing cycle, snRNAs are delivered to the assembly and undergo dramatic conformational changes through rearrangement of base pairing among themselves and with the pre-mRNA. U1 snRNP interacts with the 5’ splice site by complementary pairing between U1 snRNA and nucleotides of pre-mRNA, and U2 snRNA base-pairs with the intron to form the bulged branch site helix. Upon the arrival of the U4/U6/U5 tri-snRNP, U6 snRNA is unwound from U4 to pair with U2 and the 5’ splice site, and the displaced U4 and U1 are released (Agafonov et al. 2016). Pairing between U2-U6 snRNA results in a multi-helix complex that upon folding forms the catalytic core of the spliceosome. U5 snRNP positions both the 3’ and 5’ splice sites (Fabrizio and Abelson 1990). Thus the only snRNA directly implicated in catalysis is the U2-U6 snRNA complex (Madhani and Guthrie 1992; McPheeters and Abelson 1992; Zhang et al. 2019). Spliceosome assembly is reversible, with no single assembly step irreversible, allowing for regulation at any stage (Hoskins et al. 2011).

The spliceosome’s RNA active site is proposed to form by long-distance hydrogen bonding interactions involving the U6 snRNA AGC catalytic triad, a bulged residue in the U6 intramolecular stem loop (ISL; U74 in the human sequence), and the final two nucleotides at the 3’ end of the U6 snRNA ACAGA<u>GA</u> loop (Keating et al. 2010, Fica et al. 2013; Yan et al. 2016; Rauhut et al. 2016). However, unlike its counterpart in the self-splicing Group II intron, folding the spliceosome’s RNA active site requires assistance by NTC (Nineteen complex)-related proteins. Cwc2 (in *S. cerevisiae*), and its human equivalent, RBM22, are specifically implicated in folding activity. Although NTC proteins undergo a number of ATPase-catalyzed remodeling steps (rev. by Alves de Almeida and O’Keefe 2015), the only NTC-related proteins implicated in the folding of U2-U6 snRNA into its active conformation are RMB22 (human) and Cwc2 (yeast), as they are the only proteins shown to have a direct interaction with the U2-U6 snRNA complex (McGrail et al. 2009); Cwc2 also forms a direct noncovalent linkage with the NTC proteins *via* the WD domains of Prp19 (Vander Kooi et al. 2010). Both Cwc2 and RBM22 have been shown to form photo-induced cross-links with residues in the unpaired 5’ terminus of U6 and ISL region of their respective U6 snRNAs (Rasche et al. 2012); residues of Cwc2 in the zinc finger, RRM domain, and an interdomain connector are involved in the interaction with the RNA (Schmitzová et al. 2012; Lu et al. 2012), although precise interactions in the complex are not yet elucidated. Proximity between Cwc2 and the U6 snRNA is also observed in cryo-EM images of yeast spliceosomes in the B (Plaschka et al. 2017; Bai et al. 2018), B^act^ (Rauhut et al. 2016), B* (Wan et al. 2019), and C* (Zhang et al. 2017; Fica et al. 2017) and between RBM22 and U6 snRNA in human C* (Bertram et al. 2017) and C (Galej et al. 2016; Wan et al. 2016) stages of spliceosome assembly/activity. Splicing activity is completely inhibited by deletion of Cwc2 from yeast spliceosomes and restored by supplementation with exogenous Cwc2 (Rasche et al. 2012).

In this report, we analyze the affinity of the interaction of RBM22 with its RNA targets in both the U2-U6 snRNA complex of the major form of the spliceosome as well as its U12-U6_atac_ counterpart of the low-abundance minor variant. This minor spliceosome, responsible for the splicing of ~0.3% of introns in specific (although not fully characterized) subset of metazoan genomes, performs the same chemistry, albeit at a somewhat lesser rate than the reaction of the major spliceosome, and has the same intermediates (reviewed by Tarn and Steitz 1995; Turunen et al. 2013). However, there are alternative versions of snRNAs accomplishing the function performed by U1, U2, U4, and U6 snRNA (U5 snRNA is identical in both spliceosomes; U12 is a chimeric version of U1 and U2, and the sequences of U4_atac_ and U6_atac_ (the “atac” subscript originates from the alternative splice site sequence originally thought to associate with splicing by the minor spliceosome) compare to U4 and U6 snRNA of the major spliceosome. There is approximately ~60% sequence conservation between U12-U6_atac_ and U2-U6 snRNA complexes, and ~80% conservation is found in segments associated with formation of the catalytic core, *i.e.* the AGC triad, AGACACA loop, and ISL (Incorvaia et al. 1998; Otake et al. 2002). Structural differences include a bulged hinge and single-stranded region opposing the termini of both snRNAs in the U12-U6_atac_ complex (Figure 1B) in place of the heterogeneous multi-helix central junction observed in the protein-free human U2-U6 snRNA junction (the three-helix conformer of which is shown in Figure 1A; Zhao et al. 2013); therefore, the minor spliceosome poses a structurally simpler naturally occurring variant of the major spliceosome for study.

**Figure 1:**
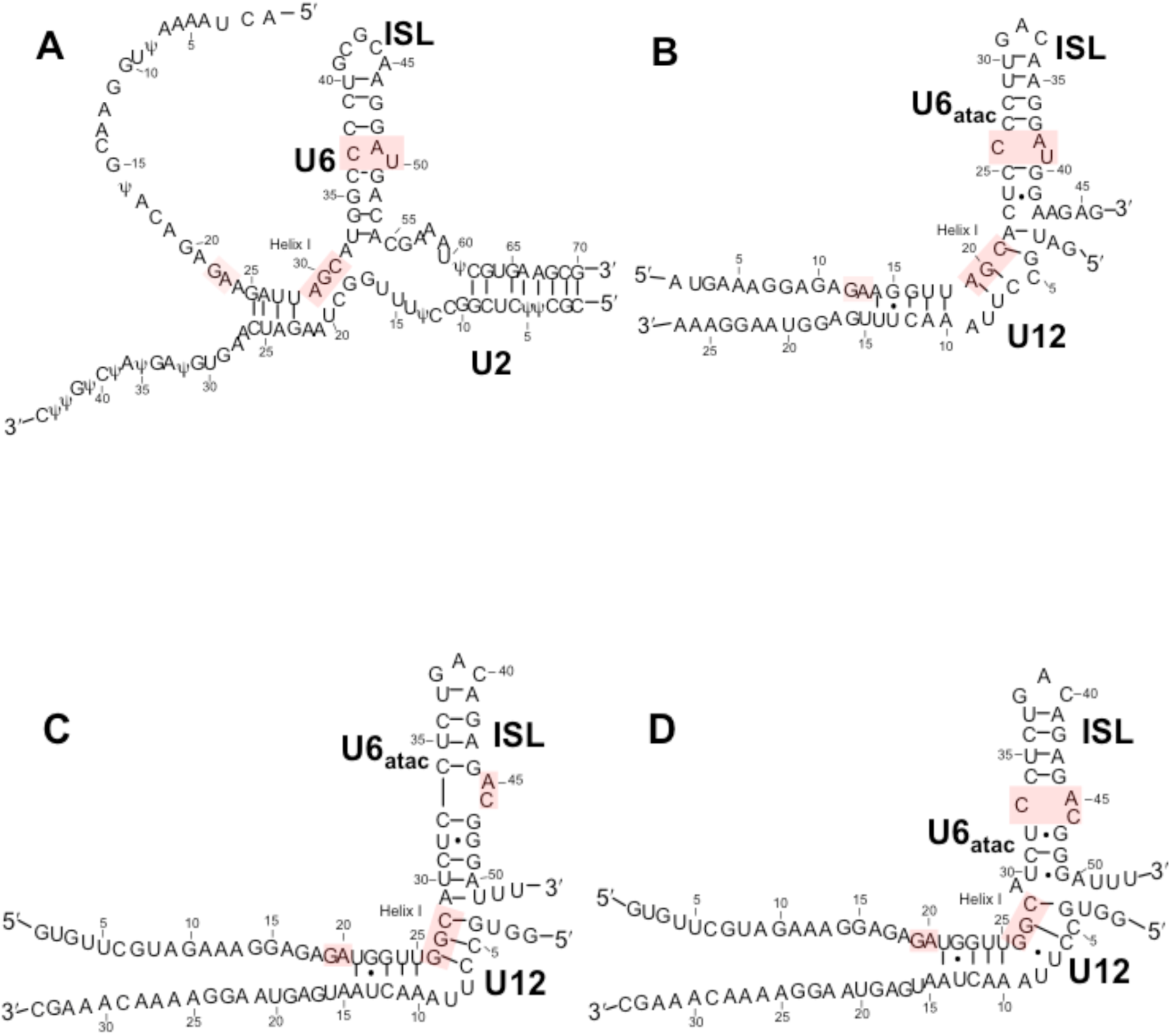
Secondary structural folds for (A) the major spliceosome U2-U6 snRNA complex in its three-helix conformation (*Note:* Ψ modifications replaced by U in transcribed samples used in these experiments); and folds proposed for: (B) the human U12-U6_atac_ snRNA complex (Tarn & Steitz 1996); and (C) the *Arabidopsis* minor spliceosome U12-U6_atac_ snRNA complex that maintains five base pairs between the catalytic triad and ISL bulge (Shukla & Padgett 1999); (D) alternate fold for the *Arabidopsis* snRNA complex (proposed here) that maintains the AY·C (where Y is a pyrimidine) motif seen in human U6 snRNA and proposed for the human U6_atac_ ISL. Lines denoting Watson-Crick base pairs and dots denoting G⋅U pairs are those proposed in the publications.

Because of the very low abundance of minor spliceosomes in metazoan cells, far less is known about the protein complement than is known for major spliceosomes. Many of the snRNP- and non-snRNP proteins are the same in both types of spliceosome or are replaced by functional analogues (Will and Lührmann 2005). However, some proteins associated with 5’ recognition and U2-branch site interaction in the major spliceosome are not represented in minor spliceosomal fractions, and several novel U11/U12-specific proteins appear to have unique functions in minor spliceosomes. While it is considered likely that the human minor spliceosome includes RBM22, its presence has not yet been documented (Will and Lührmann 2005). We ask whether RBM22 interacts with U12-U6_atac_ analogously to its interaction with U2-U6 snRNA.

In this work, we also characterize base pairing patterns of the U12-U6_atac_ complex of *Arabidopsis* and human minor spliceosomes, and verify that they adopt a pattern for the relative placement of catalytically essential elements analogous to that of key elements in the U2-U6 snRNA complex. We also analyze the affinity of RBM22 to the human U2-U6 snRNA complex by electrophoretic mobility shift assay (EMSA) and show that it binds to human U12-U6_atac_ complex with similar affinity as to U2-U6 snRNA.

Characterization of these structure-function relationships in the minor spliceosome compared to that of the U2-U6 snRNA complex provides valuable insight into the process and control of pre-mRNA splicing and provides leads to greater understanding of RNA-mediated catalysis.

## RESULTS

### Topology of protein-free human and *Arabidopsis* U12-U6_atac_ snRNA complexes

Our first goal was to characterize the base pairing pattern for the human U12-U6_atac_ snRNA complex. We started by analysis of the bimolecular construct representing the human U12-U6_atac_ snRNA complex that includes the key elements required for long range interactions associated with catalysis, *i.e.* the ACAGAGA loop, the catalytic AGC triad, and the U6_atac_ ISL (Figure 1B; sequences in SI). Analysis was performed by homonuclear and heteronuclear NMR techniques (details in Materials & Methods). We verified that the U12 and U6_atac_ snRNA strands were fully paired as a homogeneous conformation by migration of the paired complex as a single band, different from that of individual strands, on non-denaturing PAGE (Fig. S1). A 1D spectrum of imino protons illustrated 24 peaks of similar peak areas, consistent with the anticipated number of hydrogen-bonded imino protons undergoing medium-slow exchange.

1D and 2D spectra of imino/amino protons displayed line broadening that we attributed to tumbling anisotropy caused by the long single stranded 5’ region of the full U6_atac_ snRNA strand (originally included because of its equivalence to the U6 snRNA that forms photo-cross-links with the protein RBM22; Rasche et al. 2012). Subsequent NMR studies were carried out using a unimolecular construct in which the 3’ end of the truncated U6_atac_ strand was linked to the U12 snRNA strand by a stable tetraloop (Figure 2A). Folding of the unimolecular U12-U6_atac_ construct into a single conformation was also confirmed by a single band of the anticipated size in non-denaturing PAGE (Figure S1). Identical chemical shifts were shown by overlaying spectra for the two constructs in 1D and 2D NOESY and TOCSY spectra, indicating structural equivalence of the areas probed (data not shown).

**Figure 2:**
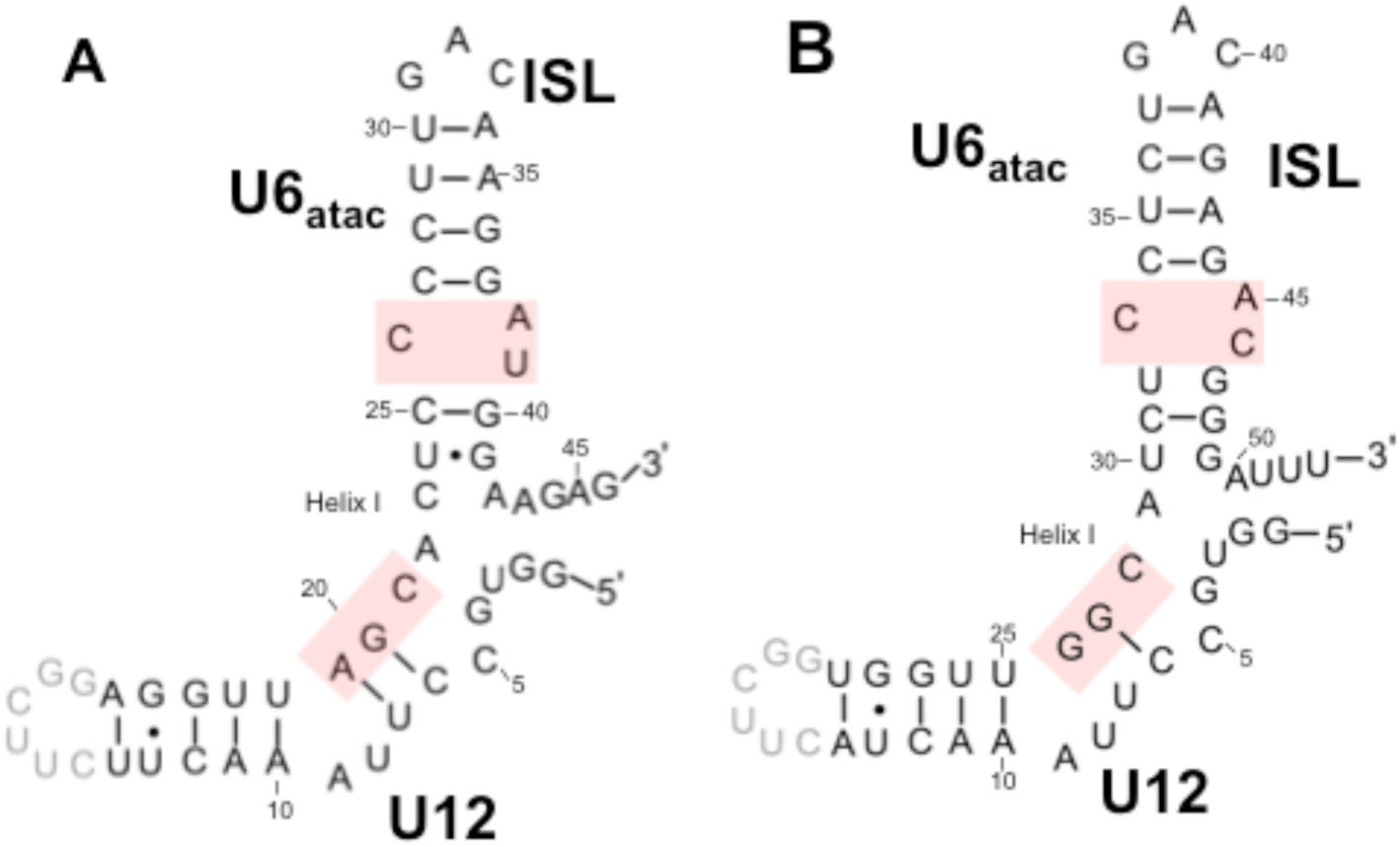
Unimolecular constructs of the (A) human and (B) *Arabidopsis* U12-U6_atac_ snRNA complexes used for NMR experiments. In each case, a stable UUCG tetraloop (in grey font) was used to connect the 3’ truncated U12 strand and the 5’ end of an abbreviated U6_atac_ strand. The catalytic triad AGC (GGC in *Arabidopsis)* and ISL bulge elements implicated in folding to form a catalytic complex (highlighted in pink, along with the final two nucleotides of the GGAGA<u>GA</u> loop not included in this construct) are preserved in the unimolecular constructs; numbering for preserved elements in corresponding snRNAs is the same as in Figure 1B and C/D. Lines denoting Watson-Crick base pairs and dots denoting G·U pairs are those identified in NMR spectra reported here.

Assignments of imino, amino, and adenine H2 protons were made from 2D NOESY spectra of the unimolecular construct (in 95%H_2_O/5%D_2_O) by analysis of intra- and inter-nucleotide nOes (Figure 3, top and center panels). Definitive identification of base pairs was aided by the distinctly different ^15^N chemical shifts of imino protons of uridine *vs*. guanosine in a 2D ^15^N-^1^H HSQC spectrum (Figure 3, bottom panel). Sequential nOes were analyzed from 2D NOESY and TOCSY spectra of non-exchangeable protons, assisted by 2D ^13^C-^1^H HSQC and 3D NOESY-HSQC experiments using a uniformly ^13^C-^15^N labeled unimolecular U12-U6_atac_ snRNA construct. Chemical shifts for aromatic, ribose (1’, 2’, and 3’), imino and amino protons are listed in Table I (SI).

**Figure 3:**
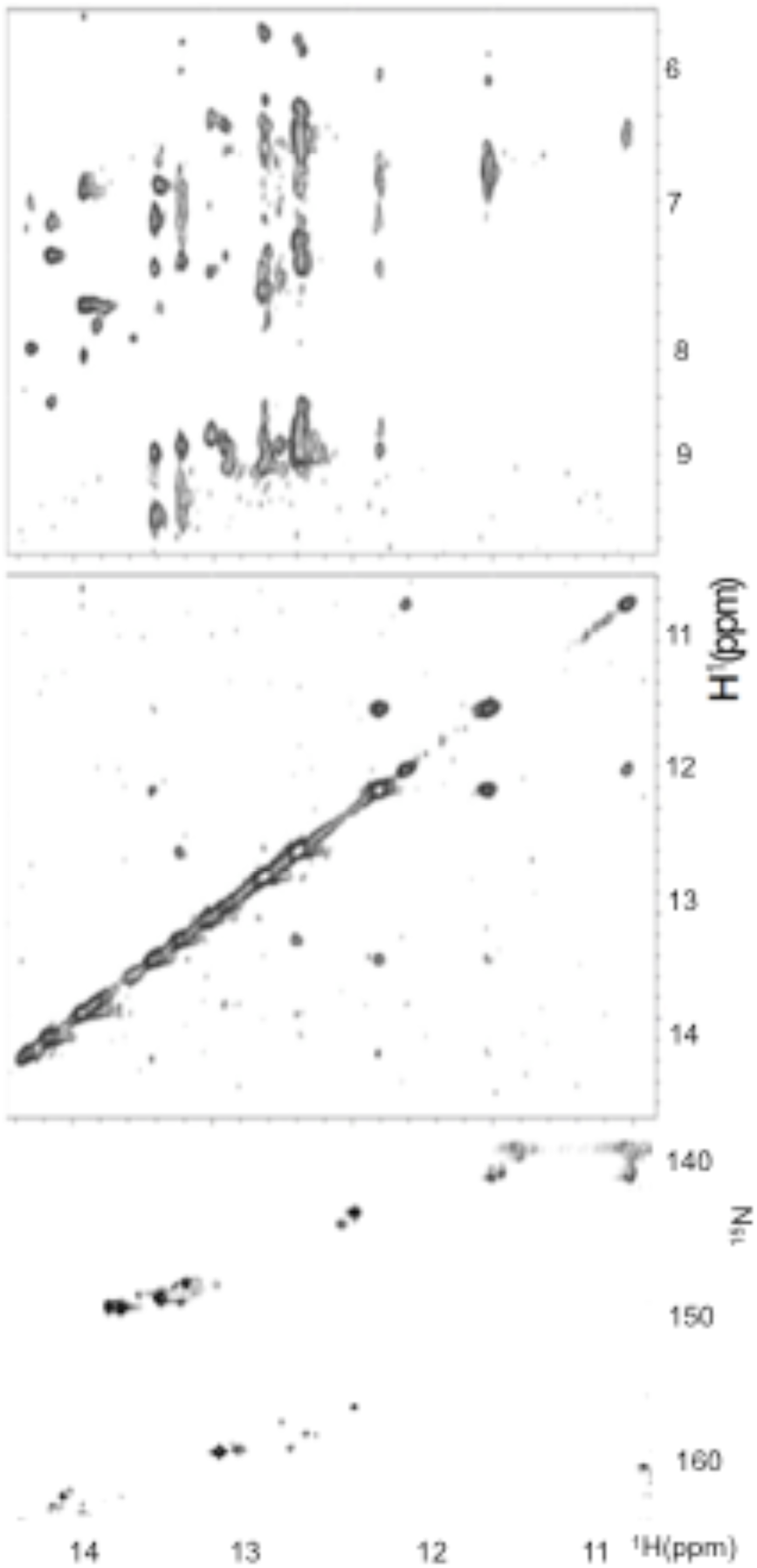
**(top panel)** Imino-amino and **(middle panel)** imino-imino regions of a NOESY spectrum of exchangeable protons of the human U12-U6_atac_ snRNA complex (Fig. 1B). Base pairing assignments from these NMR spectra confirm a base pairing pattern consistent with the fold originally proposed for the human U12-U6_atac_ snRNA complex (Tarn & Steitz 1996). Spectrum acquired with mixing time 150 ms, 10 °C, at 600 MHz. The sample was ~0.5 mM RNA in 10mM NaPi, 50 mM NaCl, 95% H_2_O, 5% D_2_O. **(bottom panel)** ^1^H-^15^N HSQC spectrum illustrating through-bond interactions between imino ^1^H and adjoining ^15^N atoms was used to confirm assignments of imino protons. The ^13^C-^15^N labeled sample was ~0.2 mM RNA in 10mM NaPi, 50 mM NaCl, 95% H_2_O, 5% D_2_O. Details of sample preparation and acquisition of NMR spectra are in Materials and Methods.

From NOESY spectra of exchangeable protons, we identified four sharp resonances of imino protons attributed to Watson-Crick base pairs and upfield shifted imino protons attributed to two G·U wobble pairs between U31-G48 and U13-G22 (Figure 2A). We assigned all five A-U, six G-C and two G·U pairs anticipated for the fold depicted in Figure 1B. Continuous A-type stacking of base pairs was observed by continuous imino-imino and aromatic-anomeric nOes between nucleotides on the 5’ side of the construct, through the ISL hairpin loop; there were breaks in continuity on the 3’ side at the point where the U6_atac_ 3’ and U12 5’ termini separate from the loop, and again at the U12 snRNA bulged (“hinge”) region. Base pair interactions defined by exchange-protected imino protons, and the proposed fold, are shown in Figure 1B. This base pairing pattern suggests a similar sequence and base pairing environment for each of the critical ion binding regions of the human U12-U6_atac_ complex as seen in the U2-U6 snRNA complex (*e.g.* Huppler et al. 2002; Sashital et al., 2004; Burke et al., 2012), and agrees with the fold originally proposed by Tarn and Steitz (1996).

To investigate the possibility of deviations from A-type helical parameters (in particular, alterations in distances or twist) in the single stranded region opposing the 3’ terminus of U6 and 5’ of U12_atac_, we measured the relative intensity of NOESY cross-peaks between protons in A-C and C-U steps of the U6_atac_ strand and compared them with cross-peaks of equivalent protons/steps in A-type duplex regions. More specifically, we compared corresponding distances between sets of protons in this single- *vs.* A-type helical double-stranded regions. In the single stranded segment, for example, we measured the relative intensity of the cross-peak of the A22H8-C23H6 step from peak volumes in a NOESY spectrum (Figure 4A), from which we calculated a distance of 3.12 Å. By comparison, the distance between A11H8-C12H6 in the A-type Helix I of the U12 snRNA segment of the unimolecular construct (Figure 2A) was 2.75 Å. Increased distance values were also reported for C23H6-U24H6 measuring 3.29 Å compared to 2.72 Å and 2.61 Å for equivalent protons in the steps C12H6-U13H6 and C28H6-U29H6 (in duplex regions) respectively. (A table of all distances is in SI). These increased distances imply that the three nucleotides in the single-stranded region undergo elongation and/or increased twist beyond that observed in A-type duplexes, which perhaps allows for increased rotation to facilitate formation of the tertiary interactions associated with formation of the active site.

**Figure 4:**
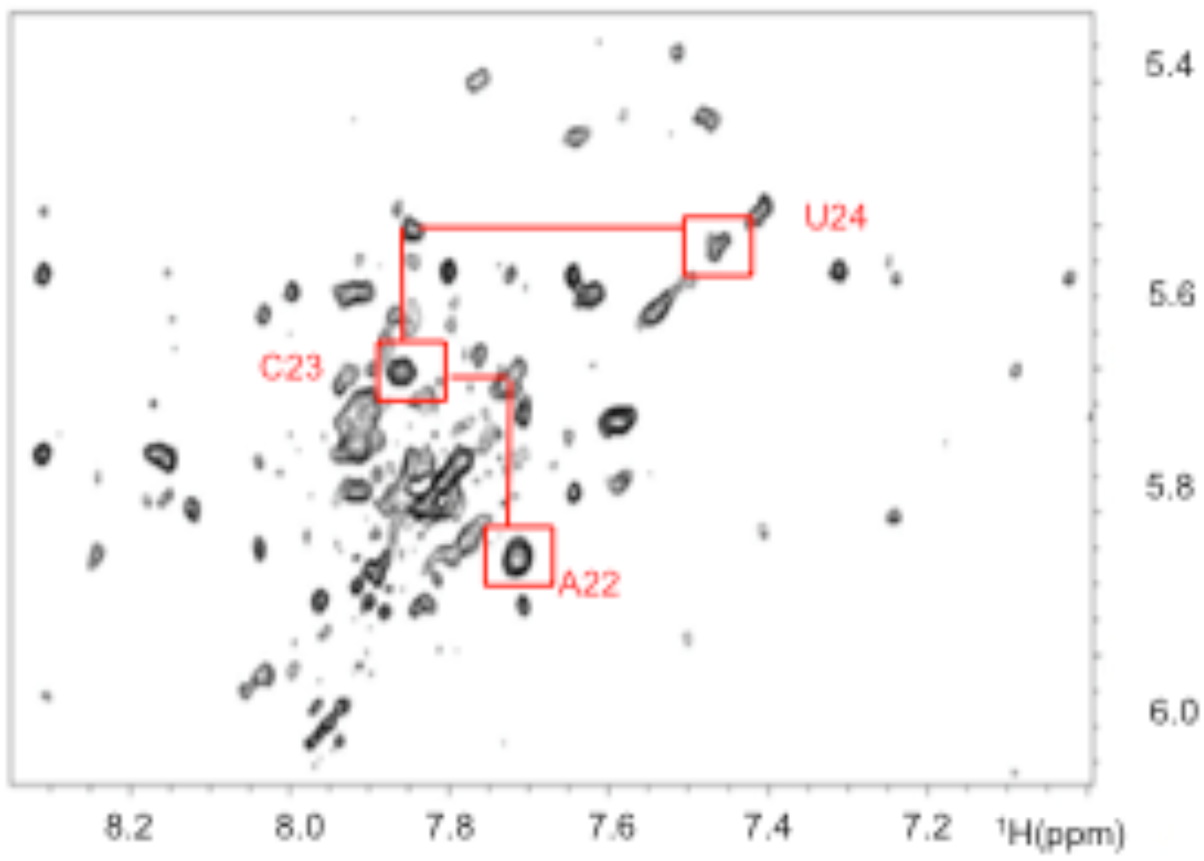
Aromatic-anomeric (base-H1’) region of a NOESY spectrum of nonexchangeable protons used for analysis of sequential assignments. Nucleotides and connectivities identified in red font identify base-H1’ nOes involving residues in the single-stranded segment of the human U6_atac_ strand opposing the 5’ and 3’ termini. Decreased intensities of intra- and inter-nucleotide base-H1’ nOes between neighboring nucleotides, relative to equivalent di-nucleotide steps in A-type helical regions, suggest that the single stranded segment opposing the 5’ and 3’ termini adopts a greater elongation between bases. Distances obtained for base-H1’ and base-base nOes in the single-stranded segment of the ISL of human and *Arabidopsis* U12-U6_atac_ constructs and corresponding steps in duplex regions are included in the text and in Table S-2 and S-3 of Supplementary Information. The sample was ~0.5 mM RNA in 10mM NaPi, 50 mM NaCl, in 99.996% D_2_O. The spectrum was collected at 600 MHz, at 25 °C, with a mixing time 250 ms. Other acquisition parameters are in Materials and Methods.

To investigate the likelihood that this perturbation of A-type helical parameters is a common feature of U12-U6_atac_ snRNA complexes, we also probed the base pairing patterns of the U12-U6_atac_ snRNA complex from *Arabadopsis.* This U12-U6_atac_ snRNA complex provided a unique opportunity because small differences in sequence may have an impact on conformation. The nucleotide sequence AUC (nt 29-31) is found in the U6_atac_ snRNA of the *Arabadopsis* snRNA complex in the single-stranded region opposing the opening, which may behave differently than ACU (nt 22-24) in the human sequence; in particular, the *Arabadopsis* U30 in the single-stranded region opposing the 3’ terminus of U6_atac_ and 5’ of U12 (unlike C23 in the equivalent position in the human sequence) allows for an additional A-U Watson-Crick base pair in the ISL, which would promote a shift in base-pairing register by one nucleotide but maintain the five base pairs as in the U2-U6 snRNA complex, as originally postulated by Shukla and Padgett (1999; Figure 1C); alternatively, the absence of the additional base pair would result in a fold with the same ISL bulge as in the human U2-U6 snRNA (Figure 1A) and the human U12-U6_atac_ snRNA (Figure 1B); this alternative fold is depicted in Figure 1D.

As for the human U12-U6_atac_ snRNA complex, we analyzed the lowest energy fold of the unimolecular *Arabadopsis* construct shown in Figure 2B by solution NMR. Folding of the unimolecular sample into a single conformation was confirmed by appearance of a single RNA band on non-denaturing PAGE. Chemical shifts in 1D and 2D spectra for equivalent base pairs were very similar to those of the human U12-U6_atac_ snRNA complex, implying that the substitution of the tetraloop to link the two strands in this construct did not perturb folding patterns.

From homonuclear NMR experiments, we assigned exchangeable and nonexchangeable protons. As in the human construct, we assigned five G-C, five A-U, and one G·U pair (there is the potential for three additional G·U pairs, but all three are exchange-broadened beyond detection). These data unequivocally fit a pattern shown in Figure 1D in which the base pairing pattern fully matches the pairing patterns expected for the fold in Figure 1B, and not for those anticipated for the fold in Figure 1C (Shukla and Padgett 1999). If the register were indeed shifted to form the pairing in Figure 1C, we would have expected to observe an additional G·U pair which, because of mid-helix position, would have been “visible”; however, we see only one G·U pair in Helix I, in the same spectral position and with the same neighbors, as found in the human complex.

We also measured distances between sets of protons in this single-stranded C28-A29 and A29-U30 steps and compared measurements with distances between corresponding protons in stacked regions. The distance for C28H6-A29H8 was 3.27 Å, compared with 2.65 Å for the same protons in the A-type helix step C40H6-A41H8 on the 3’ side of the U6_atac_ hairpin loop. For the second step, A29H8-U30H6, we measured 3.34 Å; in the absence of a duplex control of the identical sequence, we used another A-pyrimidine step, A11H8-C12H6, for which the distance was 2.68 Å. We therefore concluded that a similar pattern of internucleotide extension exists in the *Arabidopsis* U12-U6_atac_ snRNA complex as was observed in the human U12-U6_atac_ counterpart. Internucleotide base-H1’ distances for these steps followed a similar pattern, *i.e.* those in single-stranded steps indicated greater distances than their counterparts in A-type duplexes. All distance measurements are outlined in SI.

### Binding affinity of NTC-related protein RBM22 to human U2-U6 and U12-U6_atac_

To quantify and compare the binding affinity of the NTC-related protein RBM22 to RNA constructs representing the U2-U6 snRNA complex of the major spliceosome and the U12-U6_atac_ snRNA complex of the minor spliceosome, we used EMSA techniques. Starting with the bimolecular U2-U6 snRNA complex including sequence elements analogous to those in U6 snRNA shown to cross-link with RBM22 (Rasche et al. 2012; Schmitzová et al. 2012), *i.e.*, the single-stranded 5’ terminal region of U6 and the ISL hairpin loop (Figure 1A), we verified complete pairing of strands by migration of a single shifted band and disappearance of bands corresponding to individual strands on a nondenaturing gel (details in Materials and Methods). The monomer state of RBM22 was confirmed by the presence of a single band of appropriate size after electrophoresing using non-denaturing PAGE (data not shown).

For each EMSA assay, we mixed 30 μM RNA with protein concentrations ranging from 6-42 μM. After electrophoresing on a horizontal two-way nondenaturing polyacrylamide gel (details in Materials and Methods) and staining with ethidium bromide (EtBr) to visualize RNA, we observed a single band for protein-free RNA that migrated toward the anode according to construct size. Upon adding protein to the RNA sample, we observed a new band migrating at a slower rate (still toward the anode) that increased in intensity with each increase in protein concentration, and a concomitant decrease in intensity of the band representing free RNA. At a concentration ratio of 1.2:1 and 1.4:1 protein:RNA, we noted the absence of the free RNA band and the maximum intensity of the RNA-protein band, suggesting complete binding. As a control for non-specific interaction, we used a RNA fragment representing U5 snRNA (30 μM, sequence in SI) in place of U2-U6 snRNA. There was no observable shift in the protein or RNA bands upon mixing the U5 fragment with RBM22 (data not shown), consistent with specificity for the U2-U6-RBM22 interaction. Intensity of each unbound and RBM22-shifted RNA band was quantified by UVP VisionWorks software (uvp.com/visionworks) and plotted data as outlined below.

The same gel was then stained with Coomassie Blue for visualization of protein. We observed RNA-free RBM22 as a single band with minimal migration toward the cathode. For each sample of RNA (30 μM) mixed with increasing aliquots of RBM22 (6-42 μM), the band corresponding to the basic protein (only) disappeared and a band appeared on the anode side in the identical location of the shifted RNA band visualized by EtBr. Only with the final concentrations of 36 μM and 42 μM (ratios of 1.2:1 and 1.4:1 protein:RNA) did we see evidence of unbound protein. We quantified the intensity of each Coomassie-stained band with UVP VisionWorks software (uvp.com/visionworks) and plotted the data as before (Figure 6A).

**Figure 5:**
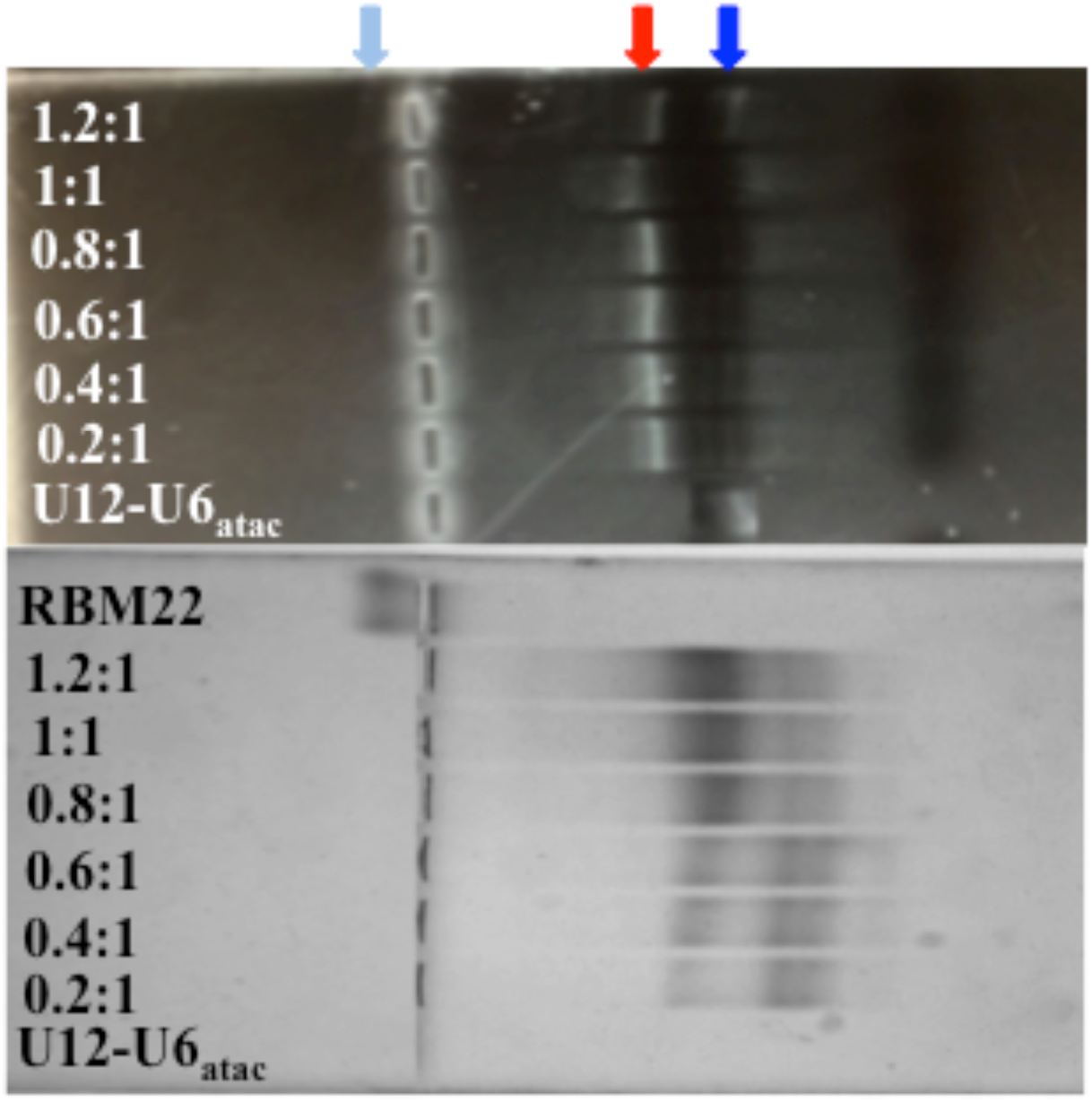
EMSA studies to quantify affinity between human RBM22 and the human U12-U6_atac_ snRNA complex (Figure 1B) were performed with 30 μM U12-U6_atac_ and concentrations of RBM22 ranging from 6μM-42μM. **(top panel)** Horizontal non-denaturing PAGE stained with EtBr visualizes bound (red arrow) and unbound RNA (blue arrow); **(bottom panel)** the same gel stained with Coomassie Brilliant Blue displays the bound protein complex (red arrow) and free protein (light blue). Further details are in Materials and Methods.

**Figure 6:**
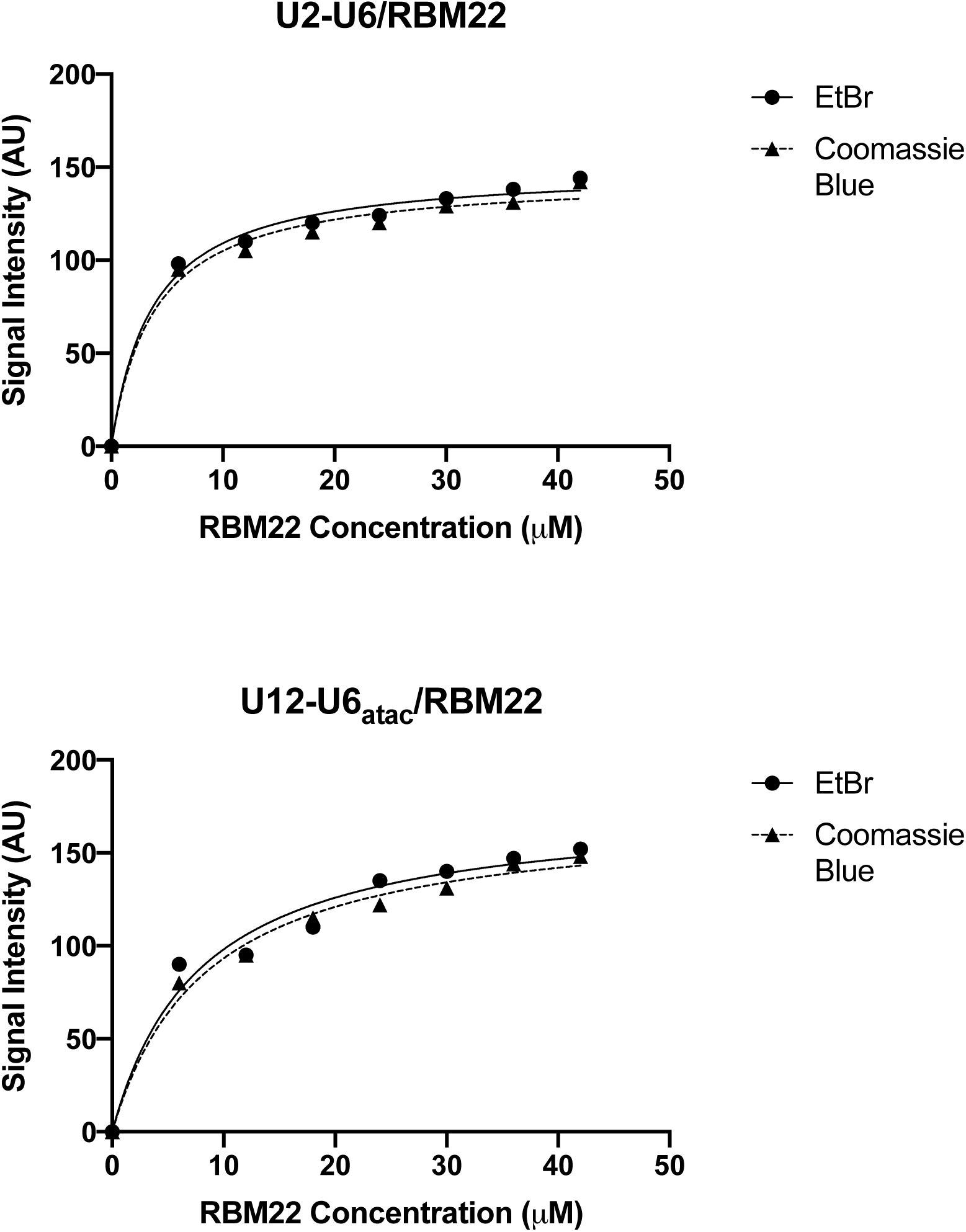
Determination of the dissociation constant K_d_ for binding of RBM22 to (A) the U2-U6 snRNA and to (B) the U12-U6_atac_ snRNA complex was performed by quantification of intensity of shifted and unshifted bands from staining by EtBr and then Coomassie Blue (see Figure 1A and 1B for constructs tested, respectively; see Figure 5 for sample gel). Concentration of protein added to 30 μM RNA was plotted *vs*. band intensity for staining with EtBr (circles) and Coomassie Blue (triangles) for each RNA complex using GraphPad Prism. The best fit for each set of points was a curve corresponding to 1:1 stoichiometry. Dissociation constants (K_d_) for binding of U2-U6 snRNA to RBM22 are 3.2±0.5 μM (EtBr)/3.8 ± 0.7 μM (Coomassie), for a mean K_d_ = 3.5 μM; for binding of U12-U6_atac_ snRNA to RBM22 are 8.4±0.7μ M (EtBr)/7.9 ± 0.7 μM (Coomassie Blue), for a mean K_d_ = 8.2 μM. Error bars indicate the S.D. for three independent experiments.

We also performed equivalent assays in reverse, *i.e.* with varying RNA concentrations while keeping the protein concentration constant; however, quantification of EtBr results was clearer with concentration of RNA kept low and constant.

For each set of quantified bands acquired by the two staining procedures, we plotted band intensity *vs*. protein concentration using GraphPad Prism (details in Materials and Methods; graphpad.com/scientific-software/prism). For the EtBr data, the data were best fit with a line describing a 1:1 stoichiometry and a K_d_ = 3.2 ± 0.5 μM (Figure 6A). For the Coomassie Blue data for the same gel, the best fit curve indicated 1:1 stoichiometry and a K_d_ = 3.8 ± 0.7 μM (also on Figure 6A). Thus, independent of staining method, the RNA-protein interaction was described by similar parameters.

We used the same EMSA protocol to test for affinity of RBM22 to the bimolecular U12-U6_atac_ snRNA complex (Figure 1B), an interaction that has not previously been shown. We observed a similar pattern of electrophoretic shifts and specificity as seen with U2-U6, that is, an increase in intensity for a shifted RNA-protein band upon increasing RBM22 concentrations, and a disappearance of the free RNA band.

Quantification of intensity and plotting for EtBr and Coomassie Blue staining were performed as above using GraphPad Prism (graphpad.com/scientific-software/prism), revealing K_d_ = 8.4 ± 0.7 μM and 7.9 ± 0.7 μM, respectively (Figure 6B). The ~38% decrease in affinity, while in the same range as that for the U2-U6 snRNA complex, suggests that the junction of the U2-U6 snRNA complex (the only region that is markedly different in sequence or topology from the central “hinged” region of the U12-U6_atac_ snRNA complex) may participate in interaction with the protein.

## DISCUSSION

We have characterized the base pairing patterns of the U12-U6_atac_ snRNA complex of both human and *Arabidopsis* minor spliceosomes, including an analysis of the single-stranded segment opposing termini of the snRNAs; we also characterized binding properties of the human NTC-related protein RBM22 to constructs representing human U12-U6_atac_, an interaction that has not previously been demonstrated, and to U2-U6 snRNA complexes from the human major spliceosome. Using bimolecular (Figure 1B) and unimolecular (Figure 2A) constructs representing the human U12-U6_atac_ snRNA complex, results acquired from homonuclear and heteronuclear NMR data confirmed the fold originally proposed by Tarn and Steitz (1996; Figure 1B). Since all evidence indicates that the minor spliceosome employs the identical chemistry as the major counterpart (Turunen et al. 2013), it is anticipated that the catalytic site would be equivalent. It is therefore significant that we have shown an equivalent base-pairing context for each of the sequence elements of the human U12-U6_atac_ complex critical for formation of the catalytic site as found in the U2-U6 snRNA complex. In addition, we have identified the correct fold (of two options) for the *Arabidopsis* U12-U6_atac_ complex, and verify that it, too, adopts a conformation with a similar context for the catalytically essential sites.

The U12-U6_atac_ snRNA complex from *Arabidopsis* shares >65% of the sequence of the human U12-U6_atac_ complex, with conservation of catalytically essential sites *i.e.* the AGC triad, AGACACA loop (GGAGAGA in *Arabidopsis*), and ISL (Shukla and Padgett 1999). Sequence differences in the U6_atac_ between the AGC triad and ISL bulge lead to the potential for two different base pairing patterns for the *Arabidopsis* U6_atac_ ISL that present a differing number of Watson-Crick base pairs between them; using a unimolecular construct representing the *Arabidopsis* U12-U6_atac_ snRNA complex (Figure 2B), analysis of homonuclear NMR data confirmed the fold illustrated in Figure 1D, supporting a conservation in the number of base pairs between the AGC triad and ISL bulge, compared to that of the human U12-U6_atac_ snRNA complex.

There are four nucleotides in the span of U6_atac_ ISL between ion-binding sites of the catalytic triad (AGC in human major and minor spliceosomes, GGC in the minor spliceosome of *Arabidopsis*) and ISL, with no more than two stacked base pairs in either human or *Arabidopsis* minor spliceosomes; by comparison, there are five nucleotides, including four base pairs, in the corresponding segment of the U6 ISL (see Figures 1A, 1B, 1D). This difference suggests the need for an alternative way to accomplish additional rotation to bring together the two ion-binding sites (along with that of the ACAGA<u>GA</u> loop or its equivalent in the U12-U6_atac_ complex), an interaction verified in yeast by genetic studies (Fica et al. 2013), by cross-linking (Anokhina et al. 2013), and now visualized in cryo-EM images (*e.g.* Yan et al. 2016), to create the active site.

We therefore analyzed the role of the single-stranded region of U6_atac_ snRNA opposing the 3’ terminus of U6_atac_ and 5’ terminus of U12 snRNAs to determine how it may contribute to flexibility of the region. By measuring internucleotide distances between pairs of protons in individual steps of the U6_atac_ strand using the relative intensity of NOESY cross-peaks in comparison with parallel measurements of equivalent protons/steps in A-type duplex regions within the same complex, we found that internucleotide distances within these single-stranded regions are elongated well beyond typical A-type helical parameters in both the human and

*Arabidopsis* U6_atac_-U12 snRNA complexes.

Either perturbation to A-type helical structure, *i.e.* additional rise or increased twist, would be consistent with the observed increase in distance between internucleotide proton pairs, and could compensate for the fewer nucleotides between the ISL bulge and the catalytic triad to favor tertiary interaction between the ISL bulge and catalytic triad (as well as the final GA nucleotides of the U6 ACAGAGA loop (GGAGAGA in *Arabidopsis*). Elongation of internucleotide distances, although relinquinshing some of the favorable stacking energies associated with an A-type helix, provides increased flexibility that may favor enhanced rotation. On the other hand, added twist, while maintaining an A-type helix, may create unfavorable steric or electrostatic clashes among stacked exocyclic groups of the nucleotides. We therefore consider it more likely that the single-stranded three-nucleotide segment adopts the more flexible elongated conformation, perhaps with added twist in its extended state. This result is important because it may help explain how, in the absence of the central junction found in the U2-U6 snRNA complex of the major spliceosome, sufficient flexibility and bending can be achieved to enable interaction between the catalytically essential elements anticipated for formation of the active site.

The most noticeable difference between the sequences of the human U2-U6 snRNA and U12-U6_atac_ snRNA complexes resides within the central region, which forms a multi-helix junction in U2-U6 complex and a simple opening in U12-U6_atac_ complex for entrance of the 5’ end of U12 and exit of the 3’ end of U6_atac_. In its *in vitro* (protein-free) state, the major spliceosome’s U2-U6 snRNA central junction is characterized by dynamic equilibrium between conformers with four and three helices (Zhao et al. 2013; Zhao et al. 2014). However, as only in the three-helix conformer observed *in situ* is the U6 snRNA AGC catalytic triad paired with U2 snRNA (as compared with pairing with the 3’ side of the U6 ISL in the four-helix conformer), a conformation that is likely to be essential for adopting the active site; while the four-helix conformer of the U2-U6 snRNA complex predominates *in vitro* (Zhao et al. 2013), a role within the assembled spliceosome is not clear. In contrast, both the human and *Arabidopsis* U12-U6_atac_ snRNA complexes adopt only a single conformer in which the catalytic triad (AGC in human, GGC in *Arabidopsis*) pairs with U12 snRNA, and therefore parallels the three-helix conformer of U2-U6 snRNA.

Binding properties of RBM22, a protein implicated in remodeling human U2-U6 snRNA prior to catalysis, to U12-U6_atac_ were analyzed by electrophoretic mobility shift assays in which we monitored migration of both protein and RNA components in the same gel. Results indicate that RBM22 binds the human U2-U6 and human U12-U6_atac_ snRNA complexes specifically with a mean K_d_ = 3.5 µM and 8.2 µM, respectively. RBM22-snRNA affinities in the same range for the two RNA complexes suggest that the protein performs the same role in both spliceosomes.

We speculate that the somewhat greater affinity for the snRNA complex of the major spliceosome results from interaction of the NTC-related protein RBM22, which is proposed to participate in remodeling or stabilizing formation of the active site, through additional or transient interaction with flexible nucleotides in the U2-U6 snRNA open three-helix junction in addition to regions in the 5’ segment of U6 snRNA and in the ISL already shown to cross-link to RBM22 (Rasche et al. 2012). If so, the absence of this junction in the human U12-U6_atac_ snRNA may help explain its somewhat lesser affinity for the protein and the slower rate of catalysis exhibited by the minor spliceosome than its major counterpart.

## MATERIALS AND METHODS

### Formation of snRNA complexes

All RNA constructs representing snRNA duplexes were transcribed from double-stranded DNA templates (synthesized by IDT, Inc.) using T7 RNA polymerase expressed and purified in the laboratory. The sequence of human U12 snRNA extends from the natural 5’ terminus to a 3’ end truncated beyond any pairing interactions with itself or with U6_atac_ snRNA. Its pairing partner, U6_atac_ snRNA, includes the full natural sequence until its truncation at A27, 13 nucleotides beyond any pairing interactions within the U6_atac_ snRNA ISL (Figure 1A). Thus upon pairing, all elements anticipated to be double stranded and/or involved in formation of a catalytic center were included in each sequence.

To minimize line broadening in NMR experiments most likely associated with the long string of single stranded nucleotides regions in the unpaired 5’ region of U6_atac_, we also designed a unimolecular U12-U6_atac_ snRNA construct that includes the key regions of the native complex in a “chimeric” construct in which the 3’ end of the truncated U6_atac_ strand was linked to the 5’ region of the U12 strand by a stable UUCG tetraloop (Figure 2A). For studies of base pairing patterns in the U12-U6_atac_ snRNA complex of the plant *Arabidopsis*, we created a similar unimolecular construct as above, from the native sequence of the *Arabidopsis* U12 and U6_atac_ sequences (Figure 2B).

For the construct representing the human U2-U6 snRNA complex, we used a U2 snRNA sequence truncated at C45 in Helix III, and the full native sequence for U6 snRNA. All sequences are specified in Supplementary Information.

Transcribed RNA was purified by denaturing PAGE; the desired band (identified by UV shadowing) was electroeluted, precipitated with ethanol, exchanged with the preferred buffer for NMR or binding experiments (conditions specified with description of experiments and in legends) using an Amicon Ultra-4 filter (MW cutoff 3 kDa; Merck Millipore, Ltd.), dried, and resuspended to the final concentration used for solution NMR studies in purified H_2_O or D_2_O, or for binding experiments in H_2_O.

For heteronuclear NMR experiments, uniformly ^13^C-^15^N-labeled samples for human unimolecular U12-U6_atac_ snRNAs were transcribed with ^13^C-^15^N-labeled NTPs (Cambridge Isotope Laboratories, Inc.) and purified as above.

To analyze pairing and folding, U12-U6_atac_ snRNA samples were heated to 70 °C for 3 min in 10 mM NaPi, 50 mM NaCl, pH 6.5, and cooled at room temperature for 30-45 minutes. Aliquots were loaded onto a 20% nondenaturing gel and electrophoresed at 100 V for 4 h at 4 °C; gels were then stained with ethidium bromide. For the bimolecular constructs, equimolar amounts of U12 and U6_atac_ strands were paired by the above protocol and pairing was verified by shifting of bands of individual strands to a single band. For both unimolecular and bimolecular constructs, formation of a homogenous fold was confirmed by appearance of a single band analyzed by nondenaturing-PAGE (Figure S1). The number of peaks observable in the imino region of 1D ^1^H NMR spectra for each snRNA construct was commensurate with all anticipated Watson-Crick and non-Watson-Crick pairs, consistent with a single conformation for each construct.

### NMR spectra

To analyze base-pairing patterns of *Arabidopsis* and human U12-U6_atac_ snRNA samples, homonuclear and heteronuclear NMR spectra were acquired on a Bruker Avance III 600 MHz spectrometer (Hunter College of CUNY, New York, NY). For examination of exchangeable protons, samples were exchanged into 10mM NaPi, pH 6.5, 50 mM NaCl and dried, and were suspended in 95% H_2_0/5% D_2_0; for spectra of nonexchangeable protons, dried samples were suspended in 99.996% D_2_0. Final RNA concentrations were in the range of 0.2-0.4 mM (concentrations of individual samples specified in figure legends) in 10 mM NaPi, 50 mM NaCl, pH 6.5.

One-dimensional spectra of exchangeable ^1^H were acquired with a 3-9-19 WATERGATE pulse sequence for water suppression. Chemical shift assignments were obtained from two-dimensional NOESY spectra of exchangeable and nonexchangeable experiments. Quadrature detection was achieved using the States-TPPI method (Marion et al. 1989). Spectra were processed and assigned using Bruker Topspin and NMRFAM Sparky software (Lee et al. 2015). Spectra were apodized using a Gaussian function, and zero filling was performed in both dimensions.

^13^C-^1^H-HSQC, ^1^H-^15^N HSQC, ^13^C-^1^H NOESY-HSQC, and HCCH-TOCSY experiments of a ^13^C-^15^N-enriched RNA sample were apodized using a Gaussian function, and zero filling was performed in both dimensions. Two-dimensional ^1^H-^15^N HSQC spectra of were acquired with a 3-9-19 WATERGATE pulse sequence for water suppression. Spectra were processed and assigned using Bruker Topspin and NMRFAM Sparky software (Lee et al. 2015).

### Expression and Purification of RBM22

*E coli* cells (Rosetta-2) were transformed with a pETM11 vector bearing the full sequence for human RBM22 protein (plasmid a gift from Prof. Reinhard Lührmann, Max Plank Institute for Biophysical Chemistry). RBM22 For protein expression, cells were grown at 37 °C in LB media containing 50 μg/mL kanamycin until 0.6 OD at 600nm was achieved. Media was adjusted with 200 μM IPTG for induction and cells were grown overnight (>16 h) at 17 °C while shaking at a rate of 225 rpm.

Purification was adapted from the protocol of Rasche et al. (2012). Cells were harvested by centrifugation at 15,000 x g during 30min and cell pellets stored at −70 °C. For purification cells pellets from 2 L of cell growth were resuspended in 30 mL IEX Buffer-A (50 mM HEPES-NaOH pH 7.5, 2 mM β-mercaptoethanol) containing 2 μL RNase I (Ambion) and lysed by sonication with a Sonic Dismembrator (Fisher Scientific) and centrifuged at 15,000 x g for 1 hour. The lysate supernatant fraction was filtered at 0.22 μm and loaded onto an ÄKTA Purifier System provided with a HiTrap SP HP 5 mL column for ion exchange chromatography. Proteins were eluted with a gradient using 50 mM HEPES-NaOH pH 7.5, 2 mM β-mercaptoethanol, 1M NaCl. Fractions containing proteins were pooled and loaded on the same system provided with a HisTrap HP 5 mL column for immobilized metal affinity chromatography (IMAC). Protein was eluted with a gradient using 50mM Tris pH 8.5, 500 mM NaCl, 1 M imidazole. Pooled fractions containing the protein were dialyzed against working buffer (20 mM phosphate buffer, pH 6.5, 100 mM NaCl, 1 mM DTT, 5% glycerol). Protein purity and integrity were tested by SDS-PAGE and migrated as a single band, with size expected for its molecular mass of 50,264 Da. The protein also ran as a single band on non-denaturing PAGE, at a migration rate consistent with monomer formation.

### Binding affinity of human U2-U6 and U12-U6_atac_ to protein RBM22

EMSA techniques were used to quantify affinity between bimolecular U2-U6 and U12-U6_atac_ RNA constructs and protein RBM22. Horizontal PAGE (5% 19:1 acrylamide:bis-acrylamide in 0.2x MOPS/histidine, pH 6.5) with wells located in the center of the gel allowed for migration anionic and cationic biomolecules or complexes to migrate towards the anode and cathode, respectively. Analysis of protein-RNA affinity was performed using 30 μM RNA human U12-U6_atac_ or human U2-U6 snRNA complexes, or nonspecific RNA, and RBM22 (6 μM – 42 μM) for ratios of 0.2:1 - 1.4:1 (total volume 15 μL per sample). Gels were electrophoresed at 100 V in a Bio-Rad Mini-Sub Cell® System, with running buffer (25 mM histidine, 30 mM MOPS, pH 6.5) at 4 °C for 1.5 h.

Gels were first stained with ethidium bromide and visualized and photographed for RNA migration by transillumination at 305 nm. Gels were then washed in 200 mL H_2_O fixed in 100 mL 10% acetic acid/50% methanol for one hour, followed by staining with 100 mL 0.05% Coomassie® Brilliant Blue R-250 in 10% acetic acid overnight, then destained with 100 mL 10% acetic acid for two hours and washed in 200 mL H_2_O overnight. Gels were then visualized and bands quantified with a UVP GelDoc-It^TM^ Imaging System equipped with a Gel HR Camera. Band intensity was plotted using GraphPad Prism® for calculation of dissociation constants (Figure 6).

## ACKNOWLEDGEMENTS

The authors thank Lenura Ziyadinova, Anjelica Gangaram, Nazir Jalili, and NamHee Kim for technical assistance with preparation of RNA and protein samples and EMSA experiments, and the NMR facility at Hunter College of CUNY and Dr. Matthew Devany for guidance with NMR experiments. NLG acknowledges financial support for this research from the PSC-CUNY grant program (award ENHC-46-84).

