## Supplementary Information for "Topology of the U12-U6atac snRNA complex of the minor spliceosome and binding by NTC-related protein RBM22"

### Nucleotide sequences for constructs.

For the bimolecular construct representing the human U12-U6$_{atac}$ snRNA complex, we used the sequence 5′- GAUGCCUUAAACUUUGAGGUAAGGAAA-3′, to represent the U12 snRNA and the sequence of, 5′- AUGAAAGGAGAGAAGGUUAGCACUCCCCUUGA-CAAGGAUGGAAGAG -3′ to represent the U6$_{atac}$ snRNA.

The unimolecular human U12-U6$_{atac}$ “chimeric” RNA had the sequence: 5′-GGUGCCUUA-AACUUCUUCGGAGGUUAGCACUCCCCUUGACAAGGAUGGAAGAG -3′.

The unimolecular *Arabidopsis* U12-U6$_{atac}$ “chimeric” RNA had the sequence: 5′-GGUGCCUUAACUACUUCGGUGGUUGGCAUCUCCUCUGACAGAGAUGGGAUUU -3′.

For the construct representing the human U2-U6 snRNA complex, we used the sequence 5′-CGCUUCUCGGCCUUUUGGCUAAGAUCUUCUCUGUAUCUGUUC-3′ to represent the U2 snRNA and 5′- GGGACUAAAAUUGGAACGUACAGAGAGAAGAUUAGCAUGGCCCCU GCGCAAGGAUGACACGAAAUUCGUGAAGCG-3′ for the U6 snRNA fragment.

### Pairing and folding of RNA samples.

**Figure S-1:** Pairing of bimolecular Equimolar quantities of individual strands were paired and incubated as described in Materials and Methods. Samples were electrophoresed on a 20% non-denaturing PAGE and stained in EtBr. Shifting of the single band of the combined strands and disappearance of the bands of individual strands implies complete pairing and formation of a single conformation. **(A)** human U2-U6 snRNA (major spliceosome; sequence above, and secondary structural scheme in Fig. 1A in text) strands: Lane 1 = U2 strand; Lane 2 = U6 strand; Lane 3: paired U2 and U6 snRNA strands.

1 2 3

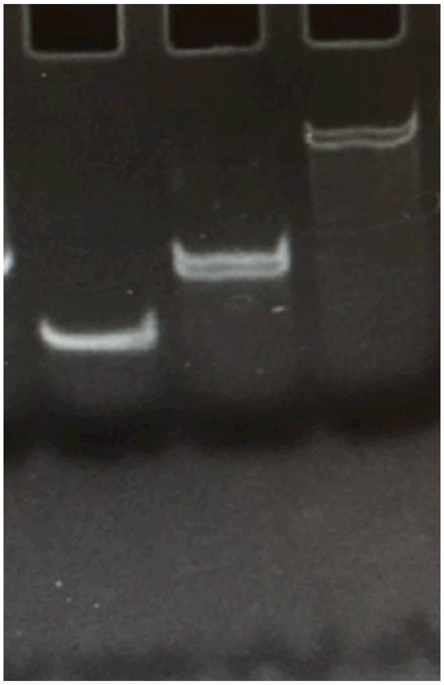

1A

**(B)** human U12-U6$_{atac}$ snRNA (minor spliceosome; sequence above, and secondary structural scheme in Fig. 1B in text) strands: Lane 1 = U12 strand; Lane 2 = U$_{atac}$ strand; Lane 3 = paired strands. See Materials and Methods (in the text) for details.

**1B**

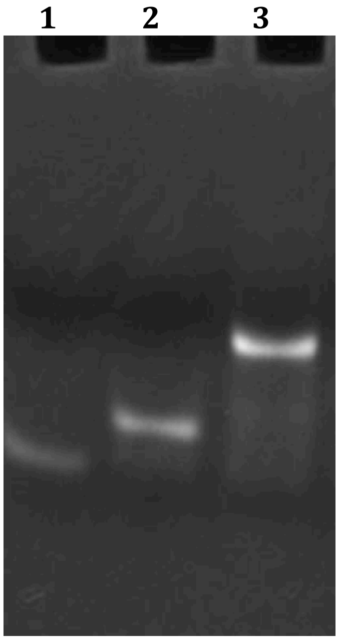


**Figure S-2A:** Folding of bimolecular (left) and unimolecular samples of the human U12-U6$_{atac}$ snRNA complex (right); sequences are above, and scheme of predicted folds in Fig. Fig. 1B and 2A, respectively). Both samples migrate as single bands, indicating homogeneity of the fold; the additional length of strands in the bimolecular construct results in retarded migration. Samples were prepared and electrophoresed on 20% non-denaturing PAGE as described in Materials and Methods, and were stained with EtBr. Appearance of a single band for each sample implies that each sample adopted a homogeneous fold. Migration was consistent with formation of the monomer for each unimolecular sample.

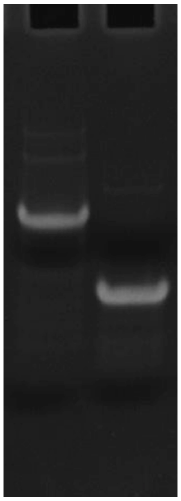

**2A**

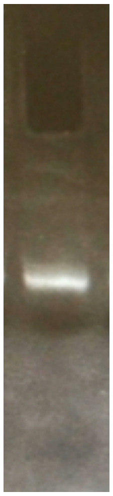

**2B**

**Figure S-2B:** Folding f the unimolecular sample *Arabidopsis* U12-U6$_{atac}$ snRNA complex (right; sequence above, and scheme of predicted fold in Fig. 2B).

**Table S-1:** NMR Chemical shifts for $^{1}H$ and $^{15}N$ (imino) in the human U12-U6$_{atac}$ snRNA unimolecular construct (Fig. 2A in text). Shifts were derived from NMR experiments described in Materials and Methods/Results in the text, and were the same for corresponding nucleotides in the bimolecular construct (Fig. 1B in text).

| Residue | H8/H6 | H5/H2 | H1′ | H2′ | H3′ | H1/H3 | $^{15}N1/^{15}N3$ |
|---|---|---|---|---|---|---|---|
| U3 | 7.75 | 4.58 | 5.41 | 4.29 | 4.51 | 13.93 | 161 |
| G4 | 7.82 | - | 5.96 | 4.71 | 4.47 | 13.2 | 143 |
| C5 | 7.78 | 5.34 | 5.60 | 4.22 | 4.44 | - | - |
| C6 | 8.03 | 5.84 | 5.71 | 4.31 | 4.37 | - | - |
| U7 | 7.70 | 4.52 | 5.38 | 4.20 | 4.44 | 13.16 | 157 |
| U8 | 7.82 | 4.99 | 5.48 | 4.41 | 4.35 | 13.81 | 158 |
| A9 | 8.36 | 7.42 | 6.07 | 4.46 | 4.88 | - | - |
| A10 | 8.24 | 7.35 | 5.98 | 4.40 | 4.80 | - | - |
| A11 | 8.16 | 7.12 | 5.75 | 4.35 | 4.75 | - | - |
| C12 | 8.07 | 5.86 | 5.74 | 4.28 | 4.32 | - | - |
| U13 | 8.04 | 5.97 | 6.13 | 4.46 | 4.29 | 10.52 | 156 |
| U14 | 8.00 | 5.90 | 6.10 | 4.38 | 4.25 | 13.96 | 161 |
| A14 | 8.09 | 7.10 | 5.87 | 4.24 | 4.60 | - | - |
| G15 | 7.13 | - | 5.76 | 4.64 | 4.57 | 11.22 | 142 |
| G16 | 8.48 | - | 5.87 | 4.18 | 5.16 | 12.42 | 148 |
| U17 | 7.85 | 4.68 | 5.56 | 4.35 | 4.57 | 14.21 | 163 |
| U18 | 7.90 | 4.72 | 5.64 | 4.44 | 4.64 | 14.62 | 164 |
| A19 | 8.32 | 7.38 | 6.11 | 4.44 | 4.85 | - | - |
| G20 | 7.88 | - | 5.78 | 4.61 | 4.72 | 13.46 | 148 |
| C21 | 7.48 | 5.29 | 5.40 | 4.51 | 4.60 | - | - |
| A22 | 7.75 | 7.04 | 5.84 | 4.10 | 4.24 | - | - |
| C23 | 7.84 | 5.33 | 5.65 | 4.25 | 4.30 | - | - |
| U24 | 7.42 | 4.51 | 5.52 | 4.28 | 4.15 | 10.2 | 153 |
| C25 | 7.67 | 5.23 | 5.54 | 4.45 | 4.47 | - | - |
| C26 | 7.73 | 5.41 | 5.56 | 4.15 | 4.31 | - | - |
| C27 | 7.69 | 5.31 | 5.55 | 4.21 | 4.45 | - | - |
| C28 | 7.75 | 5.51 | 5.92 | 4.07 | 4.16 | - | - |
| U29 | 7.83 | 4.64 | 5.52 | 4.36 | 4.22 | 13.5 | 162 |
| U30 | 7.86 | 4.70 | 5.61 | 4.53 | 4.32 | 13.8 | 165 |
| G31 | 7.76 | - | 5.88 | 4.39 | 4.69 | 12.43 | 146 |
| A32 | 8.08 | 7.05 | 5.93 | 4.22 | 4.36 | - | - |
| C33 | 7.82 | 5.62 | 5.51 | 4.31 | 4.53 | - | - |
| A34 | 8.11 | 8.08 | 5.65 | 4.28 | 4.45 | - | - |
| A35 | 8.20 | 7.30 | 5.84 | 4.38 | 4.64 | - | - |
| G36 | 6.95 | - | 5.21 | 4.21 | 4.01 | 12.54 | 147 |
| G37 | 6.82 | - | 5.32 | 4.25 | 4.08 | 12.73 | 149 |

| Residue | H8/H6 | H5/H2 | H1′ | H2′ | H3′ | H1/H3 | $^{15}$N1/$^{15}$N3 |
|---|---|---|---|---|---|---|---|
| A38 | 8.41 | 7.48 | 5.92 | 4.49 | 4.92 | - | - |
| U39 | 7.64 | 4.17 | 4.34 | 4.28 | 4.16 | 13.16 | 151 |
| G40 | 8.23 | - | 5.92 | 4.42 | 4.71 | 13.12 | 150 |
| G41 | 6.89 | - | 5.45 | 4.32 | 4.29 | 10.72 | 140 |
| A42 | 8.02 | 7.52 | 5.73 | 4.18 | 4.32 | - | - |
| A43 | 7.98 | 7.01 | 5.68 | 4.16 | 4.28 | - | - |

**Table S-2:** Distances measured for inter-nucleotide steps for nucleotides in the ISL of single-stranded vs. double-stranded regions in the unimolecular human U12-U6$_{atac}$ snRNA complex. See Fig. 2A in the text for the construct and Materials and Methods/Results for experimental details. For each dinucleotide step, interproton distances were measured for aromatic$_n$-aromatic$_{n+1}$ and anomeric$_n$-aromatic$_{n+1}$ distances.

| single-stranded nucleotide step | H6/H8-H6/H8 distance (Å) | H1′-H6/H8 distance (Å) | double-stranded (control) nucleotide step | H6/H8-H6/H8 Distance (Å) | H1′-H6/H8 distance (Å) |
|---|---|---|---|---|---|
| A22-C23 | 3.12 | 3.48 | A11-C12 | 2.75 | 2.95 |
| C23-U24 | 3.29 | 3.53 | CU12-U13/ C28-U29 | 2.72/ 2.61 | 3.01 |

**Table S-3:** Distances measured for inter-nucleotide steps for nucleotides in the ISL of single-stranded vs. double-stranded regions in the unimolecular *Arabadopsis* U12-U6$_{atac}$ snRNA complex. See Fig. 2B in the text for the construct and Materials and Methods/Results for experimental details. For each dinucleotide step, interproton distances were measured for aromatic$_n$-aromatic$_{n+1}$ and anomeric$_n$-aromatic$_{n+1}$ distances.

| single-stranded Nucleotide Step | H6/H8-H6/H8 distance (Å) | H1′-H6/H8 distance (Å) | double-stranded (control) nucleotide Step | H6/H8-H6/H8 distance (Å) | H1′-H6/H8 distance (Å) |
|---|---|---|---|---|---|
| C28-A29 | 3.27 | 3.51 | C40-A41 | 2.65 | 2.97 |
| A29-U30 | 3.34 | 3.61 | A11-C12 | 2.68 | 3.02 |
